# PCQC: Selecting optimal principal components for identifying clusters with highly imbalanced class sizes in single-cell RNA-seq data

**DOI:** 10.1101/2020.11.19.390542

**Authors:** David Burstein, John F. Fullard, Panos Roussos

**Author notes:** Correspondence: Dr. Panos Roussos; Icahn School of Medicine at Mount Sinai; 1470 Madison Ave, Fl. 9, Rm 107; New York, NY, 10029, USA.

## Abstract

**Summary:** Prior to identifying clusters in single cell gene expression experiments, selecting the top principal components is a critical step for filtering out noise in the data set. Identifying these top principal components typically focuses on the total variance explained, and principal components that explain small clusters from rare populations will not necessarily capture a large percentage of variance in the data. We present a computationally efficient alternative for identifying the optimal principal components based on the tails of the distribution of variance explained for each observation. We then evaluate the efficacy of our approach in three different single cell RNA-sequencing data sets and find that our method matches, or outperforms, other selection criteria that are typically employed in the literature.

**Availability and implementation:** pcqc is written in Python and available at github.com/RoussosLab/pcqc

## 1 Introduction

In large data sets, identifying hidden clusters of observations is especially challenging due to the prevalence of noisy features. Dimensionality reduction techniques, such as principal component analysis, circumvent this issue by forming composite measures of only the most essential features, allowing researchers to more readily discern meaningful trends within data. We typically select top features by retaining only those that capture the most variance across all observations. This approach, however, limits our ability to derive information from less abundant functional clusters, such as rare populations of cells associated with specific disease [19]. We propose a new methodology to better capture small clusters by selecting candidate principal components based on the distribution of the variance explained across all observations. *Existing Methodologies*. Since each principal component captures a certain percentage of the variation, we can graphically represent this relationship in the form of a scree plot. To decide the cut-off point, we look for the location of the elbow in the plot, where the amount of information captured by the principal component appears to level off. Given their conceptual simplicity and computational efficiency, scree plots are commonly used for identifying the top principal components in single-cell RNA sequencing data [5, 13]. However, visually determining this cut-off is often challenging as there can be many potential cut-off points. Alternatively, we can assess the significance of the principal components through a permutation test, where we permute the entries in the columns of our data matrix to construct a randomized data set and then evaluate whether a given principal component in the original data set captures more variance than its analogue in the permuted data. Permutation tests require significantly more computational resources than their scree plot counterparts. In addition, ranking the top principal components based on total variance captured runs the risk of ignoring principal components that play an important role in describing data points that belong to smaller clusters.

### PCQC

To address the aforementioned issue, we propose the PCQC (Principal Component Quantile Check) criteria, a computationally efficient methodology, where we rank the principal components by extracting a value near the tail of the distribution of variance explained across all observations, as opposed to the total variance explained (**Figure 1a,1b**). Intuitively, while principal components describing small clusters may not explain much variance in the data as a whole, we anticipate that for a small subgroup of data points, the variance explained is very large. As such, important principal components may be identified by focusing on observations where the principal component captures a larger percentage of variance. The toy example in **Figure 1c** illustrates the efficacy of the PCQC methodology; retaining the top two features with the largest mean or total variance explained yields only two distinct clusters, while PCQC preserves all four clusters found in the three-dimensional space. (See Supplementary Materials for details.)

**Figure 1:**
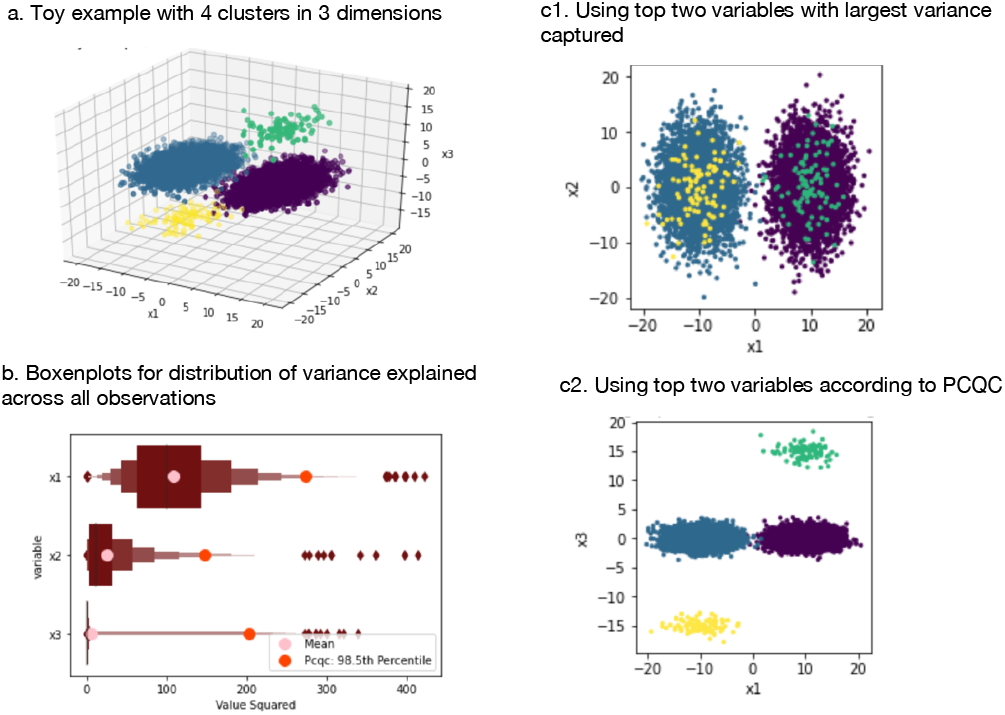
A toy example illustrating the PCQC variable selection process.

### 2 Data Sets

#### To assess the performance of the PCQC method, we consider three different single-cell RNA sequencing data sets

We used existing and in-house generated multimodal single cell gene expression data to validate the PCQC method. Fresh PBMCs were subjected to Cellular Indexing of Transcriptomes and Epitopes by Sequencing (CITE-seq) (PMID: 28759029) with single-cell libraries generated using the 10x platform (10x Genomics). This approach permits study of the transciptomes of individual cells while gaining quantitative and qualitative information on a panel of cell surface proteins (CD11b, CD11c, CD123, CD14, CD16, CD27, CD3, CD4, CD40, CD45, CD45RA, CD56, CD8 and HLA_DR) through the use of barcoded antibodies. Additionally, we assess the efficacy of our approach on a CBMC data set produced using CITE-seq and a PBMC data set generated through single-cell RNA-seq [2, 7]. We construct our ground truth labels, by clustering the smaller dimensional antibody derived tag (ADT) data from our CITE-seq experiments or by leveraging marker genes to label different cell types based on prior studies. To better understand how dimensionality reduction impacts downstream clustering analysis, we also generate a bootstraped variation of the dataset, where we re-sample the data based on empirically observed distributions of frequencies of different cell types. We preprocessed data based on common practices and filtered out doublets or low expression single cell barcodes [3, 5, 8, 18]. Based on the recommended procedure from the Seurat and Scanpy pipelines, we cluster the transcript count data by embedding the data in a k-nearest neighbor graph and extract the hidden clusters using a Louvain or Leiden community detection algorithm. Subsequently, we train a multinomial Bayes model to map the unsupervised clustering to the ground-truth labels.

### 3 Results

From **Table 1**, we observe that PCQC consistently matches, or outperforms, alternative approaches, including Scree Plots and Permutation Tests. For example, when classifying the rare dendritic cell type, the PCQC approach decreases the median log-loss score by more than 13%. **Table 1** also illustrates how selecting an arbitrary number (50 or 70) of principal components captures a significant amount of noise and results in undesirably large values for the log loss. We note that while the percentile cutoff parameter in the PCQC methodology influences the quality of downstream clustering analysis, through a priori inference, the practitioner can make informed estimations for the size of the smaller clusters and choose the appropriate percentile cut-off to yield the desired result. Given PCQC’s strong performance relative to competing methodologies, we anticipate the the PCQC approach will provide much needed perspective in flagging critical principal components for clustering data with imbalanced class sizes.

| Data Set | Pcqc | Scree | Permutation | Top 50/70 PCs |
| --- | --- | --- | --- | --- |
| 1. 8.6k PBMC Citeseq [7] | **0.87** | 0.96 | 0.88 | 0.95 |
| 2. 2.7k PBMC [2] | **0.88** | 0.91 | 0.91 | 1.04 |
| 3a. 2.7k PBMC Resampled | **0.87** | 0.91 | 0.91 | 0.94 |
| 3b. 2.7k PBMC Resampled CD14 | **0.78** | 0.89 | 0.89 | 0.81 |
| 3c. 2.7k PBMC Resampled Dendritic | **1.11** | 1.28 | 1.28 | 1.21 |
| 4a. 17k PBMC | **1.13** | **1.13** | 1.14 | 1.26 |
| 4b. 17k PBMC Doublets | **0.59** | **0.59** | **0.59** | 0.60 |

## Supporting information

Supplementary Results

## Acknowledgements

This work was supported by the National Institutes of Health (R01AG050986, R01MH109677, R01MH109897, U01MH116442, R01MH110921, R01AG065582 and R01AG067025) and the Veterans Affairs (Merit grant BX002395 Roussos). Further, this work was supported in part through the computational resources and staff expertise provided by Scientific Computing at the Icahn School of Medicine at Mount Sinai. We thank Dr. Kathryn Twyman for her insightful comments in reviewing a draft of this manuscript. The funders had no role in the design and conduct of the study; collection, management, analysis, and interpretation of the data; preparation, review, or approval of the manuscript; and decision to submit the manuscript for publication.

## Notes

### Competing Interest Statement

The authors have declared no competing interest.

