## Supplementary Results for "PCQC: Selecting optimal principal components for identifying clusters with highly imbalanced class sizes in single-cell RNA-seq data"

### Supplementary Materials

### 1. Other Methodologies and Related Work

Although the focus of our work serves to evaluate the methodologies typically employed for performing dimensionality reduction to cluster single cell RNA sequencing data, we also wish to highlight some of the less prominent approaches for reducing the number of features in the data.

In contrast to scree plots and permutation tests, information criteria, such as Bayesian Information Criteria, provide a definitive cut-off point by identifying the principal components that capture noisy data through probabilistic modeling [15]. Unfortunately, while the underlying assumptions appear relatively innocuous, modeling the principal component weights as independent identically distributed normal random variables, appears to contradict the notion that principal component weights depend on its corresponding cluster and that clusters can have heterogeneous sizes.

Alternatively, since we are primarily interested in selecting principal components for the purpose of clustering, we can also identify interesting principal components by performing a Kruskal-Wallis test to see if there are meaningful differences in the principal components in question between different clusters. While we can integrate this approach as a quick check to verify the significance of our principal components, iteratively selecting the top principal components based on a threshold for the p-values poses challenges, similar to conducting a permutation test.

The Jackstraw method, like the permutation test, identifies significant principal components by identifying statistically significant differences between the permuted and original variables in the data set and flagging PC's that capture these statistically significant variables [7]. And while the Jackstraw method has been popularized in the literature for clustering single cell RNA-seq data, the approach faces two significant drawbacks both in terms of computational cost and by thresholding principal components through p-values [1, 4, 6].

Also noteworthy, Markos et al. [14] provides an R-package for simultaneously selecting the optimal principal components and computing clusters using K-means. Nevertheless, while it has the advantage of combining two optimization procedures that are potentially at odds with one another, this approach currently only accommodates the K-means clustering algorithm.

### 2. Mathematical Motivation for PCQC

In order to flag principal components that capture significant information pertaining to smaller clusters, we need a computationally efficient approach to compute the $100 * (1 - \alpha)$th percentile of the variance explained by each principal component. Fortunately, as described below, we can compute the distribution of variance explained by using the eigenvectors (principal component) and eigenvalues by tweaking the steps taken in principal component analysis.

First, consider a data matrix $A \in \mathbb{R}^{n\times p}$, with n observations and p features. Assume, without loss of generality, that we have already transformed our data matrix such that all features have 0 mean and unit variance. We can then compute our covariance matrix as $\frac{1}{n-1}A^TA$. Observe that the ith entry on the diagonal of the covariance matrix equals the variance of the ith feature $X_i$ due to the fact that,

$$(\frac{1}{n-1}A^TA)_{ii} = \frac{1}{n-1}\sum_{k=1}^{n} a_{ki}^2 - 0 = Var(X_i)$$

Consequently, it follows that the trace, or the sum of the diagonal elements of our covariance matrix equals the total variance across all of the features,

$$tr(\frac{1}{n-1}A^TA) := \sum_{i=1}^{p}(\frac{1}{n-1}A^TA)_{ii} = \sum_{i=1}^{p} Var(X_i). \text{ (1)}$$

We can leverage this prior result to ascertain how much variance the principal components capture. Consider the singular value decomposition of $A$, such that

$$\frac{1}{\sqrt{n-1}}A = U\Sigma V^T,$$

where $U$ and $V^T$ are orthonormal matrices ($V^{-1} = V^T$) and $\Sigma$ is a diagonal matrix. We can then employ the singular value decomposition of $A$, to compute the eigenvalues/eigenvectors of the covariance matrix,

$$\frac{1}{n-1}A^TA = (U\Sigma V^T)^T(U\Sigma V^T) = V\Sigma U^TU\Sigma V^T = V\Sigma^2V^T,$$

where $\Sigma^2 := \Lambda$$ is the diagonal matrix consisting of the eigenvalues $\lambda$ and the orthonormal matrix $V$ consists of the principal components. Since the trace operation is commutative, that is for any two matrices $tr(YZ) = tr(ZY)$, we then have that

$$tr(\frac{1}{n-1}A^TA) = tr(V\Lambda V^T) = tr(V^TV\Lambda) = tr(\Lambda) = \sum_i \lambda_i$$

Based on equation (1), $tr(\frac{1}{n-1}A^TA)$ also equals the total variance across all features. We then reduce the dimensionality of our matrix by identifying the principal components that capture the most variance in the data to attain the best possible approximation of our original data matrix.

After determining the optimal number of principal components to retain, we can then reduce the dimensionality of our original matrix $A$, by taking the matrix product $AV_*$, where $V_* \in \mathbb{R}^{p \times k}$, projects our data matrix on the subspace consisting of the top $k$ principal components.

As mentioned previously, we want to be cautious about using $\lambda$ as a criterion for selecting the top principal components since it acts as an aggregate measure of variance captured across all data points and may overlook information pertaining to smaller clusters. Unfortunately, while the covariance matrix $\frac{1}{n-1}A^T A$ is helpful for understanding relationships between features, it does not provide the granularity we need for analyzing how certain principal components describe different data points. To circumvent this issue, first observe that due to the commutative properties of the trace operator,

$$tr(\frac{1}{n-1}A^T A) = tr(\frac{1}{n-1}AA^T) = \frac{1}{n-1}\sum_{i=1}^{n}\sum_{k=1}^{p} a_{ik}^2$$

At this juncture we can consider a particular observation i and measure its contribution, $\sum_{k=1}^{p} a_{ik}^2$, to the total variance measured across all features. In order to gauge the amount of variance explained by a principal component for a particular observation, we return to the singular value decomposition of the matrix. Note that since $\frac{1}{\sqrt{n-1}}A = U\sqrt{\Lambda}V^T, \Rightarrow \frac{1}{n-1}AA^T = U\Lambda U^T$, we have that

$$\frac{1}{n-1}\sum_{k=1}^{n} a_{ik}^2 = \sum_{k=1}^{n} \lambda_k u_{ik}^2,$$

by inspecting the ith diagonal entry of $\frac{1}{n-1}AA^T$ and $U\Lambda U^T$. Consequently, we can define the distribution of variance explained across all observations for principal component k, using the fact that the amount of variance explained by principal component k for observation i equals

$$\lambda_k u_{ik}^2. \quad (2)$$

Note that we anticipate that the PCQC approach will retain many of the strengths of the more traditional scree plot approach as well, since the eigenvalues still play an important role in ranking principal components, as evidenced in expression (2). And while expression (2) provides us with a deeper understanding of the PCQC methodology, in practice we can compute the distribution of variance explained in a more direct fashion. Let $A_* := AV_* \in \mathbb{R}^{n \times k}$ be the dimensionally reduced representation of our data matrix. We can then compute the contribution of the jth feature to the total variance in the reduced data matrix as,

$\sum_{i=1}^{n} a_{*ij}^2$ , and analyze the distribution of the variance explained by looking at the individual terms in the summation.

### 3. Results

After performing the preprocessing steps, we can now leverage principal component analysis to filter out noisy covariates in our cord blood mononuclear data set. As evidenced by Figure 1, using visualization techniques like scree plots present certain limitations, specifically in regards to deciding how many principal components to retain. Among the top 50 principal components, we might be inclined to choose 7 or possibly 10 principal components, by looking at potential discontinuities in the scree plot, to assess significance. However, when we look at a zoomed-in plot consisting of principal components 10-50, we see discontinuities at 10, 13 and 16.

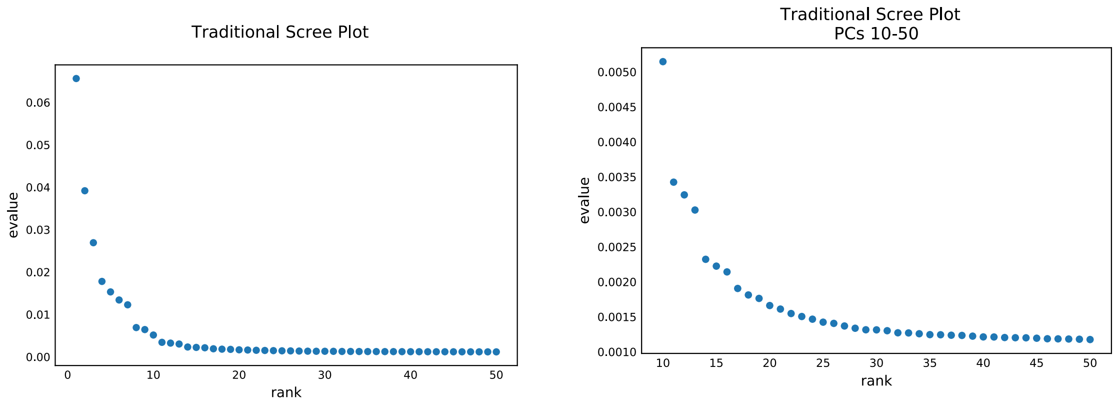


**Figure 1:** Scree plot for the top 50 principal components in the cord blood mononuclear dataset (left) and for principal components 10-50 (right)

To help resolve some of the ambiguities by visually inspecting scree plots, we also provide tables to identify significant principal components, by computing the ratio of the variance explained of two successive principal components. In particular, if a given principal component is not significant, we anticipate the ratio to be close to 1, as the variance explained should achieve an asymptotic value and the denominator acts as a proxy for a null value, similar to the permutation tests discussed later. Conversely, if there is a discontinuity in the scree plot, we would anticipate the ratio to be substantially larger than 1. As a rule of thumb, we typically look at principal components that explain 10% more variance than their successor, but other thresholds are certainly possible. By looking at Table 1, we can readily identify the principal components flagged as significant in the scree plots, without looking at different x-axis windows in the scree plot. Specifically, principal components 7,10,13 and 16 all explain 10% or more variance than their successor and since principal component (PC) 16 is the lowest ranked principal component that is also practically significant, we should consider retaining only the top 16 PCs. (In contrast, the more lowly-ranked PC 19 only captures 6% more variance than its successor and is most likely not regarded as significant.). While the table representation serves as a tool to objectively identify significant PCs, visual inspection of the scree plots could still capture useful information hidden by the table and, hence, selecting the top PCs should leverage both approaches.

**Table 1:** Scree plot significance table

| Principal Component Rank | Ratio |
|---|---|
| 7 | 1.78 |
| 1 | 1.68 |
| 3 | 1.51 |
| 10 | 1.50 |
| 2 | 1.46 |
| 13 | 1.30 |
| 9 | 1.25 |
| 4 | 1.16 |
| 5 | 1.14 |
| 16 | 1.12 |
| 6 | 1.09 |
| 8 | 1.07 |
| 12 | 1.07 |
| 19 | 1.06 |
| 11 | 1.06 |

After selecting the top 16 principal components, we can now proceed to cluster our data using the Leiden community detection algorithm. To do so, we first need to embed our data as a k-nearest neighbors graph, by choosing a value for k. When running the Leiden algorithm, we also need to decide on a value for the resolution limit $\gamma$ , which impacts the size of the clusters [10]. In our numerical experiments, we consider a range of values for these parameters, $\gamma \in \{.4, .8, 1.2, 1.6\}$ and $k \in \{10, 15, 20, 25\}$. To find a suitable clustering of our data, we perform 20 trials to account for a range of randomized starting points for each possible parameter combination.

Unfortunately, when training a Multinomial Bayes model to map the output clusters from the Leiden algorithm to the ground truth labels, the model performs sub-optimally. As illustrated in Figure 2, the median log-loss for selecting the top 16 PC's is around 0.95 but the median log-loss for the PCQC-methodology, discussed later, is only 0.87. Furthermore, by inspecting the performance of the model across individual clusters we notice that the median log-loss in classifying the ground truth CD8 cluster with the top 16 PC's is 1.4, in contrast to the 0.72 log-loss achieved by the PCQC-methodology.

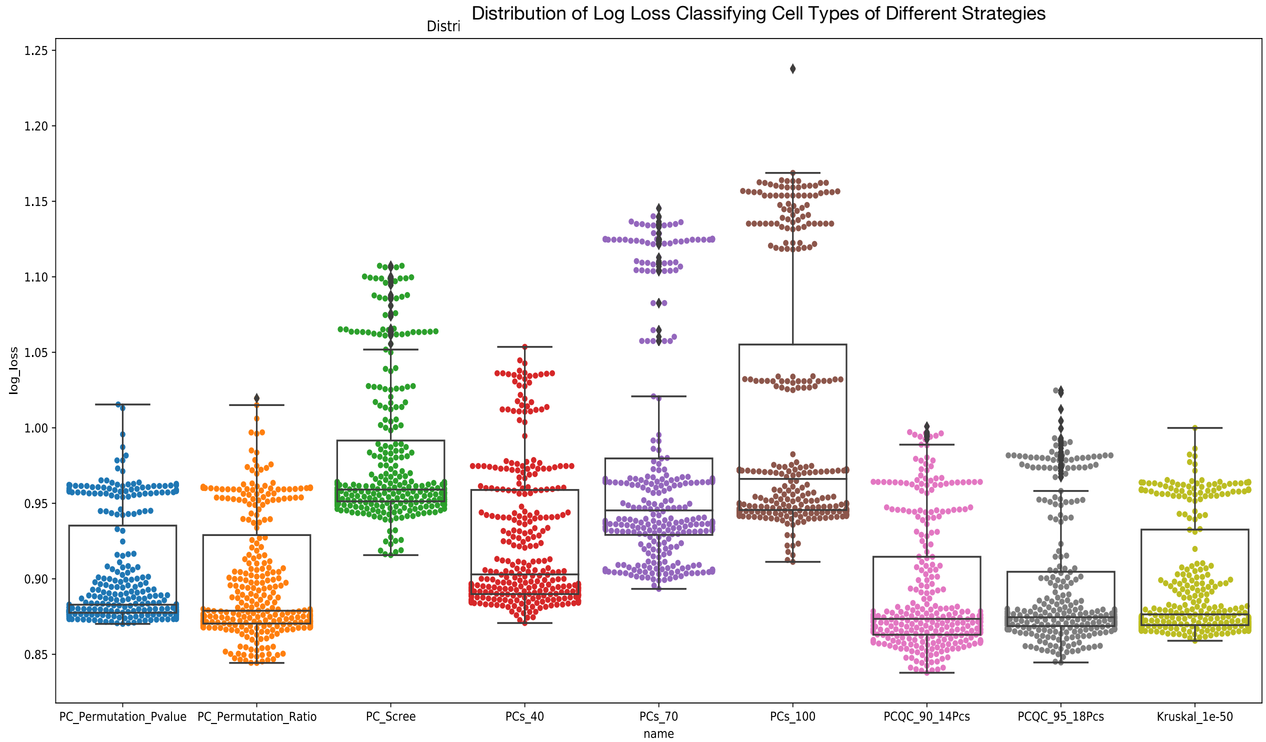


**Figure 2:** Distribution of log-loss for mapping leiden clusters to ground truth labels using a multinomial Bayes model. On the x-axis, note that PCQC_90_14Pcs indicates the performance of the clustering when selecting the top 14 principal components using the PCQC methodology with the $90^{th}$ percentile as a cutoff parameter.

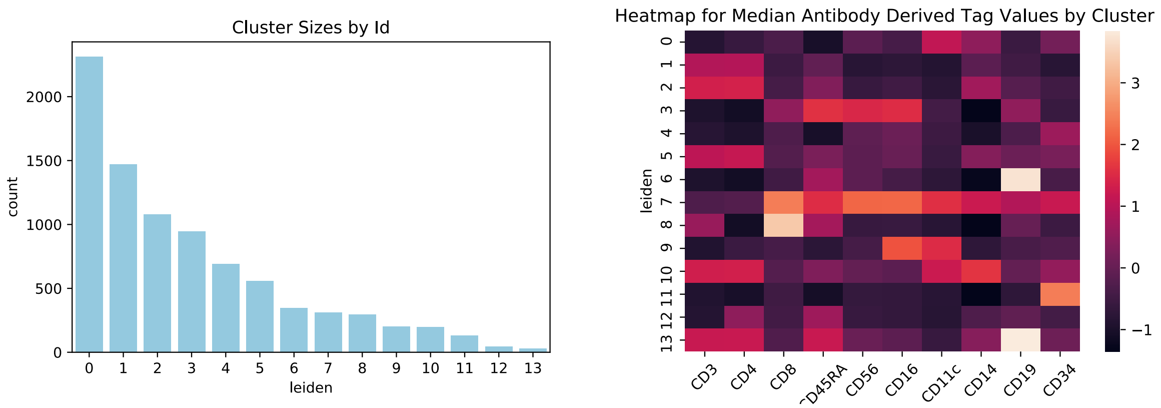


**Figure 3:** A barplot indicating the frequency for each cluster labeled from 0 to 13 (left) and a heat map indicating antibody derived tag values for each cluster (right).

To better understand why selecting the top 16 PCs by variance explained yielded such poor performance in classifying the CD8 cluster, in Figure 3 we provide summary statistics regarding each ground truth cluster. In particular, when we cluster the antibody derived tag data to form the ground truth labels, we note that cluster 8 attains high values for the ’CD8’, but this ’CD8’ cluster size is in fact relatively small, as it consists of about 300 observations out of 8,600 records. Since only about 3% of observations fall into the CD8 cluster, by focusing only on the principal components that capture the most variance on average, we have in fact discarded the principal components that describe this relatively rare cluster. At the same time, we also cannot simply choose an arbitrary number of principal components, like 40 or 100, as we would be retaining a significant amount of noise in the data, as evidenced by the high log-loss scores seen in Figure 2.

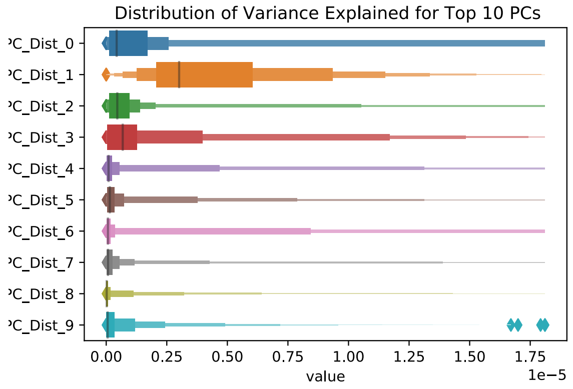


**Figure 4:** A boxenplot for the distribution of variance explained over different observations for the top 10 principal components [12]. Note the principal components are labeled from 0-9.

To address this issue, we propose the PCQC-methodology, where we select principal components based on the tails of the distribution of variance explained for each principal component. In Figure 4, we can more readily identify a glaring limitation with the traditional principal component selection process. By flagging important PCs by the total variance explained, we are in effect assessing the importance of a principal component by focusing on the mean variance explained across all data points, when in actuality, the mean is just one summary statistic describing how a particular principal component explains the variance in the data. Specifically, Figure 1 demonstrates that while PC 0 has a much larger mean than PC 1, Figure 4 tells us that the median for PC 1 is much larger than PC 0. Furthermore, we also see that certain principal components, like PC 0 and PC 6 have particularly large tails. Consequently, to identify important principal components in our data with the PCQC methodology, we rank each PC by the tails of the distribution for the variance explained, and by doing so, we can include PCs that

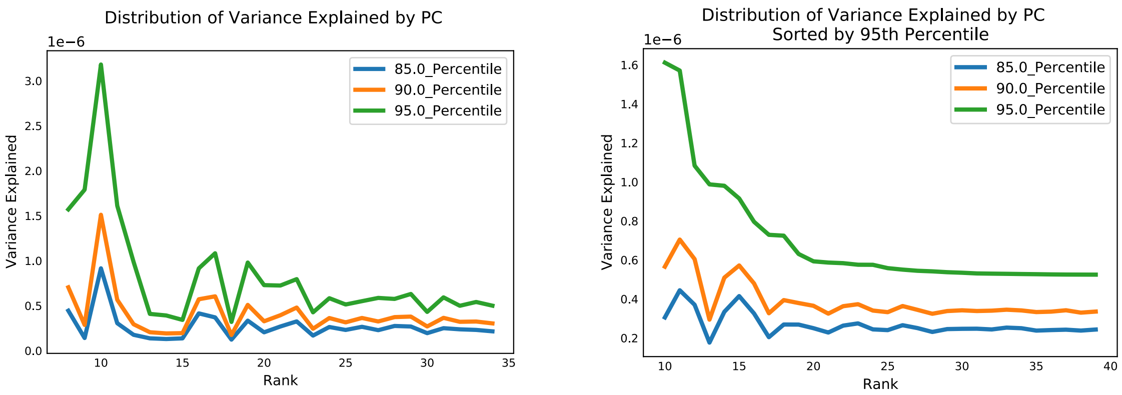


**Figure 5:** Flagging important principal components by looking at the tails of the distribution of variance explained with the PCQC methodology. Note that the PC's on the x-axis are ranked according to their mean (left). A scree plot where the principal components are ranked according to the 95$^{th}$ percentile of the distribution of variance explained (right).

describe small clusters. In contrast, if we look solely at the mean, we could end up disregarding principal components that correspond to these small clusters as they aren't representative of a typical observation in the data.

When applying the PCQC methodology to determine which principal components are in fact significant, in Figure 5, we observe that many of the significant principal components from the original scree plot are significant using the PCQC approach, as a large mean is often, but not always, indicative of a heavy tail. Nevertheless, the PCQC methodology distinguishes PCs, like PC 22, that, despite explaining a large amount of variance for a relatively small number of observations, were not flagged in the original scree plots. After computing the distribution of variance explained for each PC, we can then extract how large these tails are by looking at a particular percentile and construct a modified scree plot to determine potentially significant principal components.

We stress that, while in this particular example we observe relatively modest differences when selecting different percentile values for identifying the tail of the distribution, in general, we should consider percentiles that reflect the possible sizes of the small clusters in the data. In so doing, we generate a statistic that more faithfully captures information from these small groups. In contrast to the traditional scree plot methodology discussed above, which achieves a log loss of 1.4, by leveraging a summary statistic that can more readily capture information regarding these smaller clusters, the log loss for the CD8 cluster using the PCQC methodology is only

0.72. Furthermore, as illustrated in Figure 2, the PCQC methodology also outperforms the traditional scree plot methodology in the log-loss measure aggregated across all cell types.

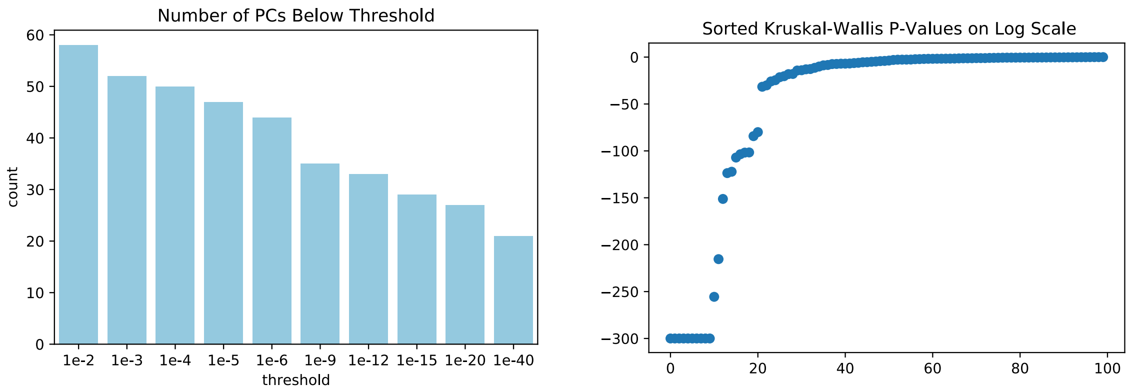


**Figure 6:** Frequency of the number of principal components with a p-value less than the threshold labeled on the x-axis (left). P-values for the top 100PCs plotted in log-scale (right).

In addition to the PCQC methodology, we also considered an alternative principal component selection criteria, where in an unsupervised fashion, we iteratively cluster our data and only retain the principal components that vary significantly between the clusters. To determine significance, we use a variation of the Krusal-Wallis test and correct for imbalanced cluster sizes by resampling the data such that when performing the Kruskal-Wallis test, the class sizes are indeed balanced. Interpreting significance using p-values is especially tricky, considering the large number of principal components with impressively small p-values. Note that in Figure 6, over 40 principal components have p-values smaller than $10^{-6}$ when, in contrast, the PCQC approach identifies less than 20 significant PCs. And even though certain cutoffs for the Kruskal-Wallis test do yield strong performance for classifying clusters, the test provides little insight as to where we should establish such cutoffs. Consequently, we recommend this procedure as a check to ensure that principal components capture meaningful differences between clusters, but not as a standalone principal component selection criterion.

In particular, we observe that the permutation test identifies 24 practically significant principal components by computing the analogous measure for significance as we performed for the traditional scree plots in Table 1, where the denominator is no longer the variance explained by the following principal component, but rather the variance explained from the corresponding principal component in the permuted data matrix. Since the permutation test assesses significance based on the mean variance explained from a given principal component, we readily

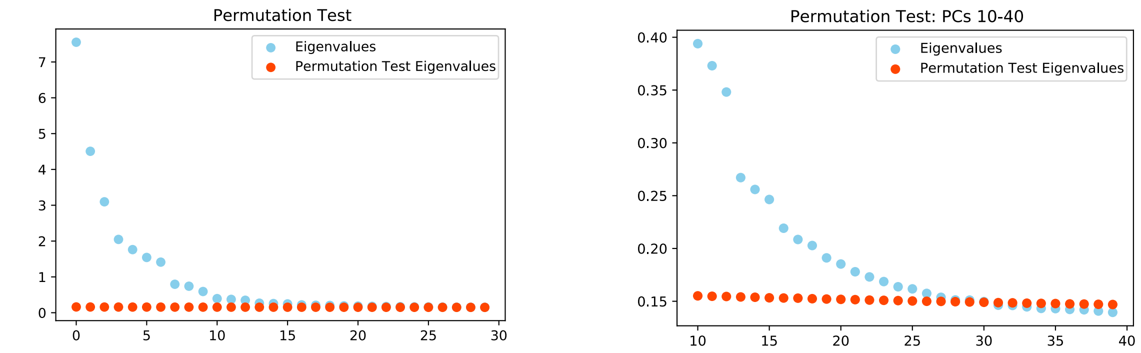


**Figure 7:** Assessing significance of principal components using the permutation test (left). A restricted window of the plot on the left focusing on PCs 10-40 (right).

note (and observe in the following examples) that the permutation test may overlook principal components that play important roles in describing smaller clusters. We also emphasize that using a p-value threshold from the permutation test also faces serious limitations, as that approach does not provide clear intuition regarding meaningful cutoff points, similar to what we encountered when leveraging the Kruskal-Wallis test.

To better understand the proposed PCQC methodology we next look at a both a peripheral blood mononuclear cell dataset containing roughly 2700 entries, as well as a resampled version of the data set where the more uncommon dendritic cells appear at a biologically plausible lower frequency. See Figure 8 for details.

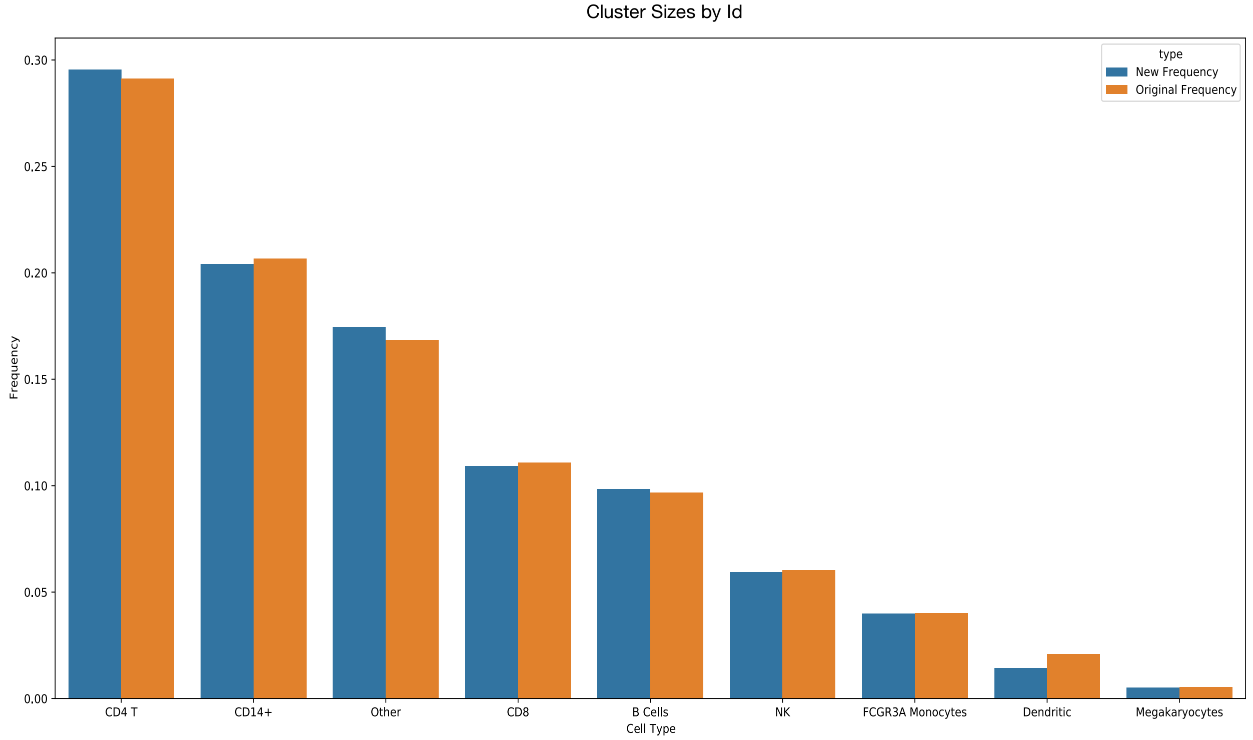


**Figure 8:** Frequencies for different cell types in the original peripheral blood mononuclear cell data set with 2700 observations and the new frequencies when performing a weighted sampling without replacement with 2300 observations, where we decrease the likelihood of observing dendritic cells.

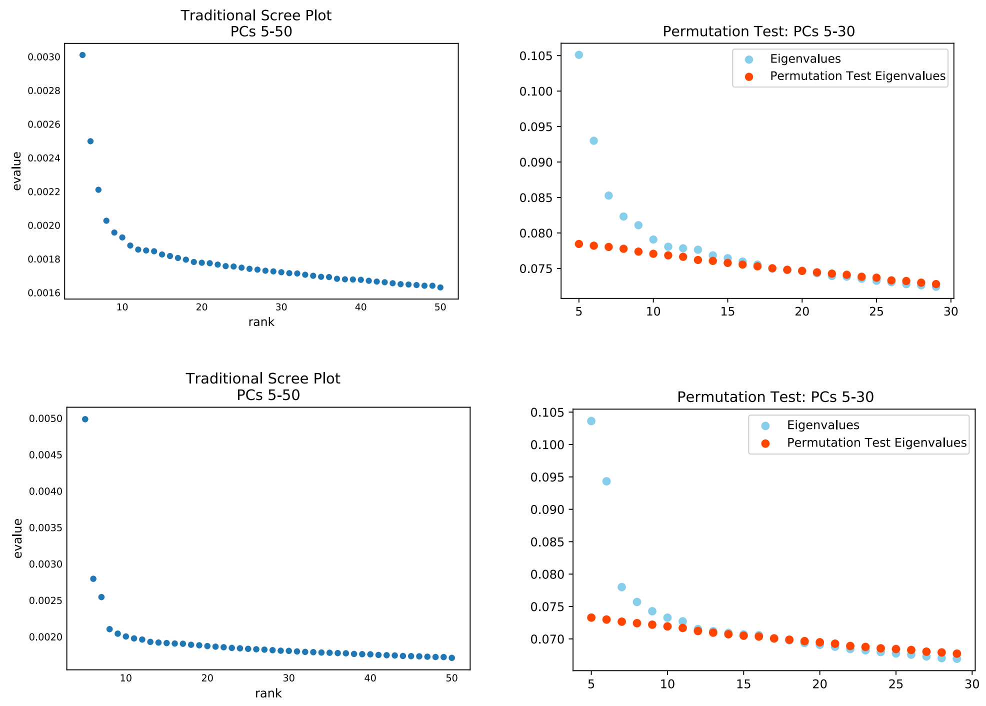


**Figure 9:** Scree plot for ranking the top principal components in the original data set (top left) and thee permutation test (top right). Both methodologies highlight 7 significant principal components using the 10% increase threshold discussed earlier. The bottom plots are the analogous ranking criteria for the resampled data.

When selecting important principal components to retain for further analysis, both the traditional scree plot and the permutation test yield 7 principal components in both the original and resampled data set, as referenced in Figure 9. In contrast, the PCQC methodology identifies 11 and 10 important principal components to retain for downstream analysis, where we rank principal components using the 99.5th percentile, as illustrated in Figure 10.

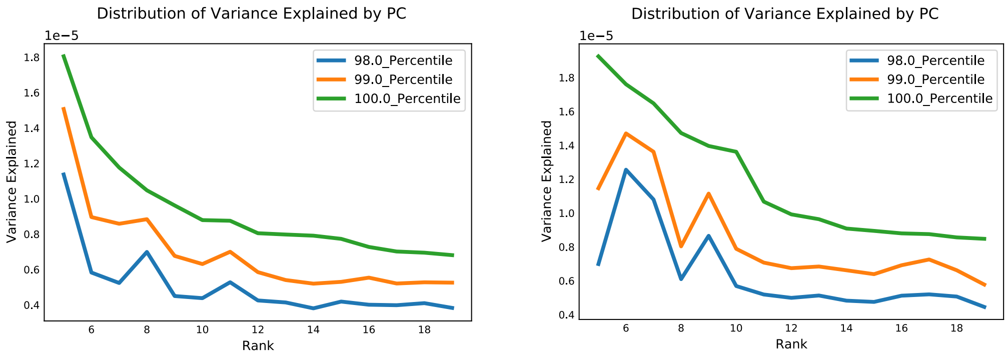


**Figure 10:** PCQC ranking for the principal components in the original data set (left) and the PCQC ranking for the resampled data set (right).

Even though we observe only modest differences between the methodologies in the original data, we do notice a more prominent distinction in log loss for the resampled data, as indicated in Figure 11. In particular, while the overall improvement in log loss appears relatively modest, when we analyze performance on individual ground truth cluster identities, we see in Figure 12 that the 99.5th percentile PCQC methodology substantially improves the clustering for the Dendritic and CD14 cell types, while retaining competitive performance on the remaining categories. Furthermore, as indicated in Figure 13, the log-loss distribution for the PCQC methodology is more tightly concentrated around its median than the alternative scree and permutation test methodologies. And, although the 98th percentile PCQC cut-off yield sub-par performance relative to the scree/permutation methodologies, this decrease in performance is not entirely surprising either. Dendritic cells account for less than 2% of the dataset and we would need to choose a higher threshold for the tail cutoff parameter to ensure that our ranking methodology captures the behavior of this rare cell type. Nevertheless, the PCQC methodology provides the practitioner with the flexibility for defining the tail of the distribution for variance explained with regards to each principal component, and through this approach, we can more readily identify principal components that play an important role for capturing variation in rare cell types, like dendritic cells.

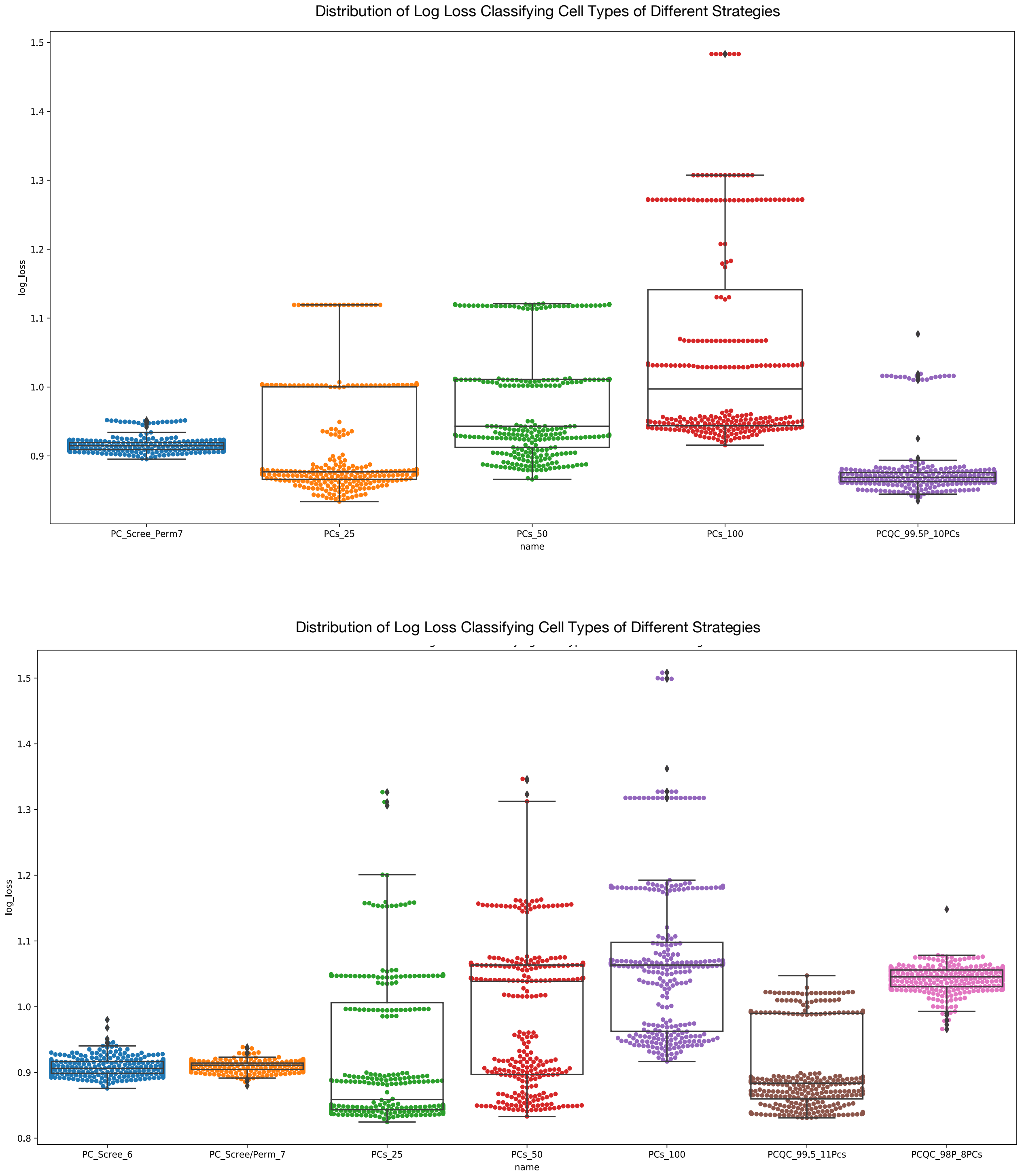


**Figure 11:** Log loss for classifying different cell types in the original PBMC data set with 2700 observations (top) and in the resampled data set (bottom).

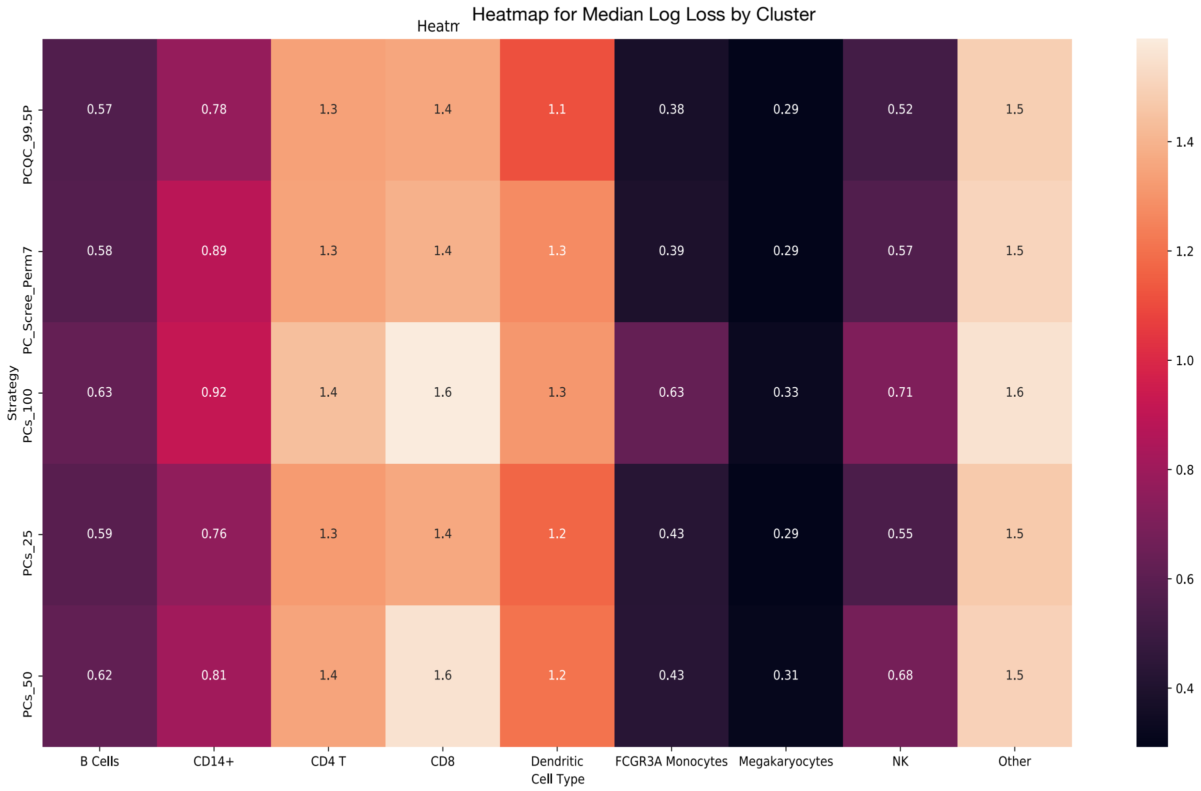


**Figure 12:** Heatmap for the median log-loss for each ground truth cluster identity across different PC selection strategies.

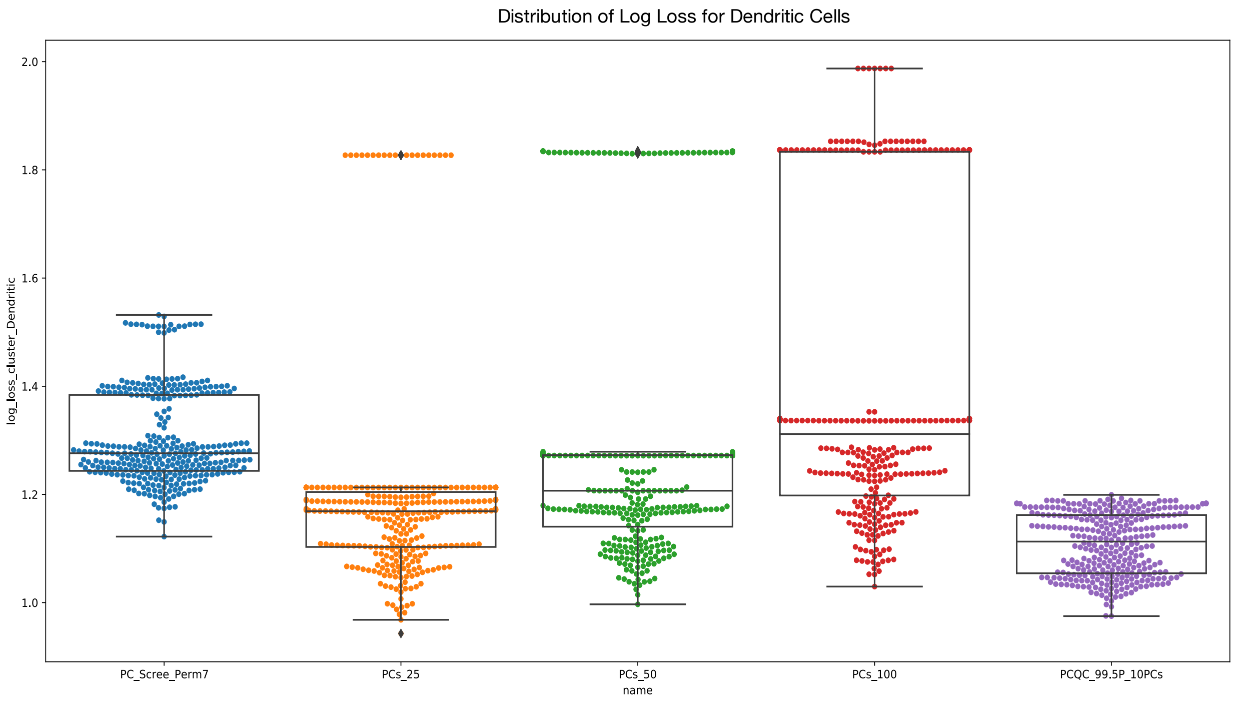


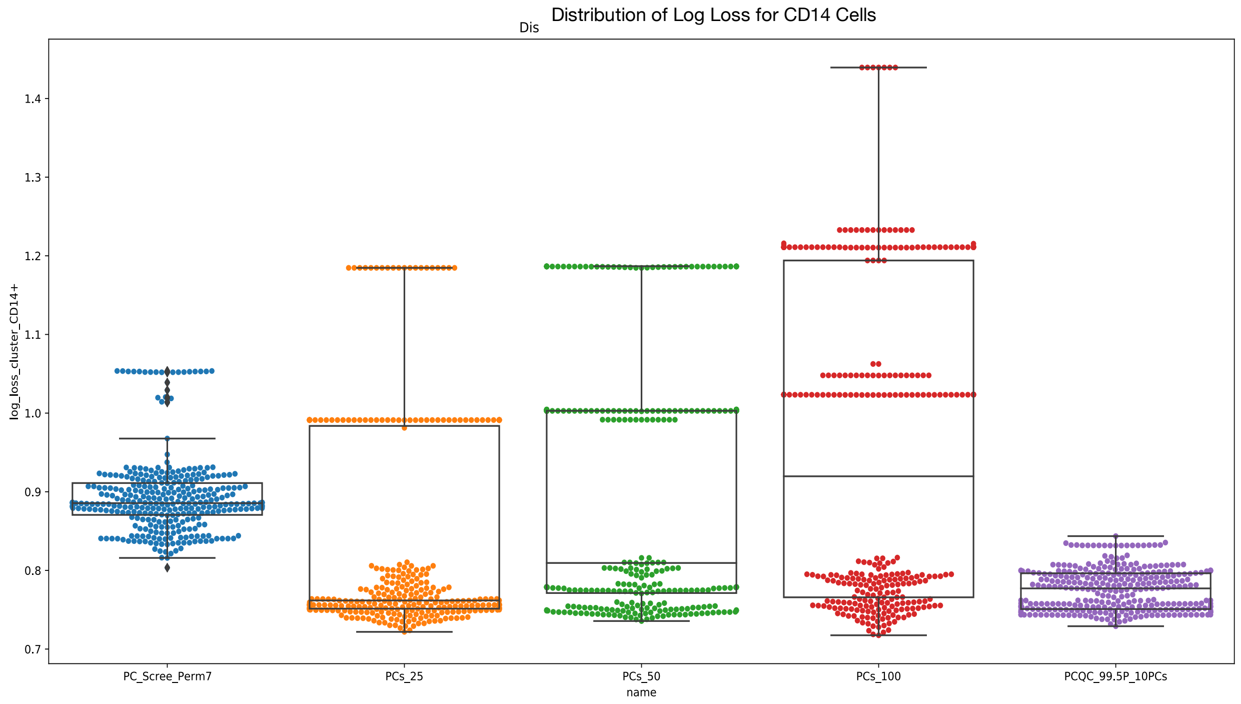


**Figure 13:** A close up view of log-loss performance for the CD14 and Dendritic cell types in the resampled PBMC 2700 dataset.

Finally, we consider a peripheral blood mononuclear cell transcript dataset with over 17k observations generated using the CITE-seq approach. We compare the performance of our proposed PCQC methodology with ground truth labels derived from the corresponding antibody derived tags and cell hashing data. Foremost, when comparing the different methodologies, while Figure 14 highlights how the PCQC methodology identifies different subsets of important principal components than the traditional scree plot, Figure 15 demonstrates the robustness of the log loss distribution from evaluating the ground truth labels across the scree plot, permutation test and 98 percentile PCQC methodologies. In contrast, Figure 15 also illustrates how selecting an arbitrary number (50 or 70) of principal components captures a significant amount of noise and results in an undesirably large value for the log loss. Furthermore, selecting an incorrect value for the percentile, can also yield suboptimal results. Subsequently in Figure 16, we confirm that a small cluster drives the high log loss score for the 95 percentile PCQC methodology.

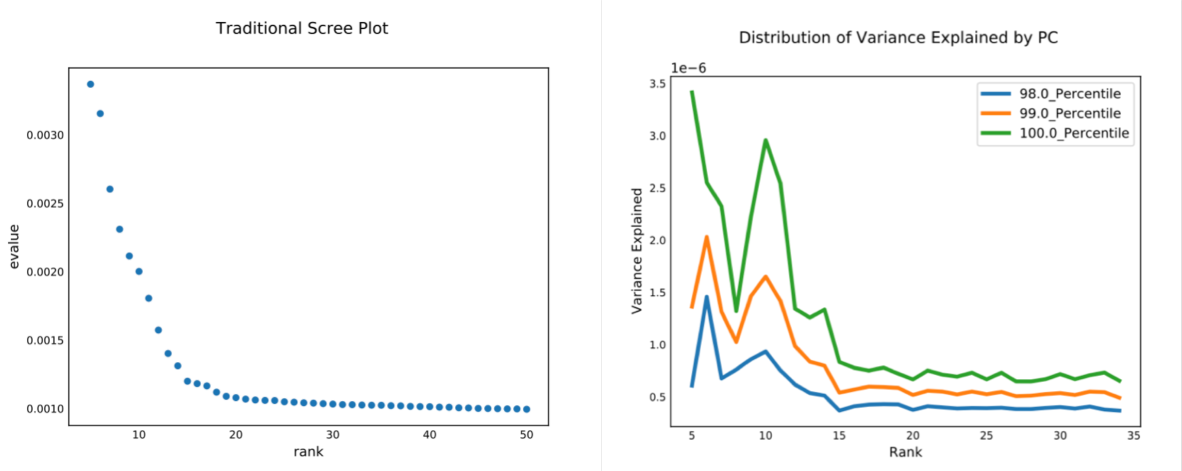


**Figure 14:** A scree plot for the 7k+ peripheral blood mononuclear cell data set (left) and the corresponding plot for the PCQC methodology (right). Note that the PCQC methodology draws greater attention to the 14$^{th}$ principal component than in the traditional scree plot. Not shown, the permutation test flags 17 principal components as significant.

At this juncture, we wish to emphasize that although at first glance the 95th percentile PCQC methodology appears inferior to the competing methodologies, we stress that we would anticipate improved performance assessing the 95th percentile PCQC methodology on a different ground truth data set where the ground truth labels are more balanced. Consequently, we create ground truth labels based on the cell hashing data, labeling approximately 80% of the cells as singlets and the remaining cells as doublets. We then evaluate the performance for these different approaches in Figure 17. In particular, we observe that the 95 percentile PCQC methodology modestly outperforms the competing methodologies by attaining a more concentrated log loss distribution.

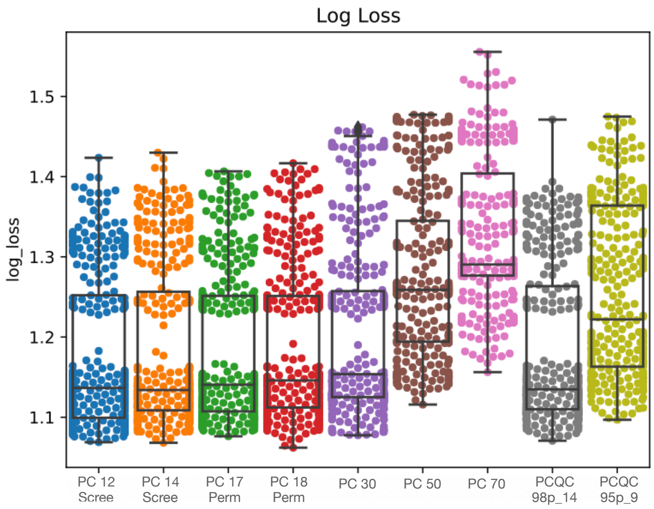


**Figure 15:** Distribution of log loss for the 17k+ peripheral blood mononuclear cell data set.

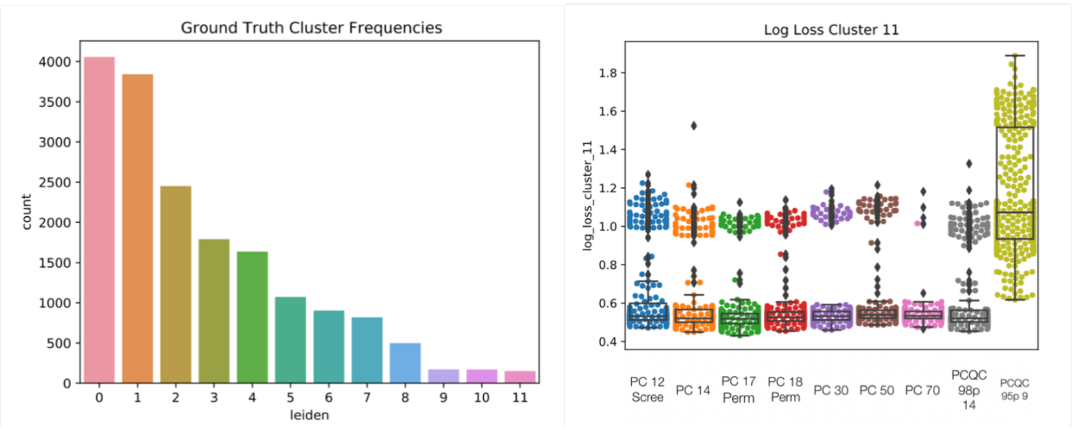


**Figure 16:** Frequencies for the ground truth cluster labels (left) and the log loss distribution for the smallest ground truth cluster, cluster 11 (right).

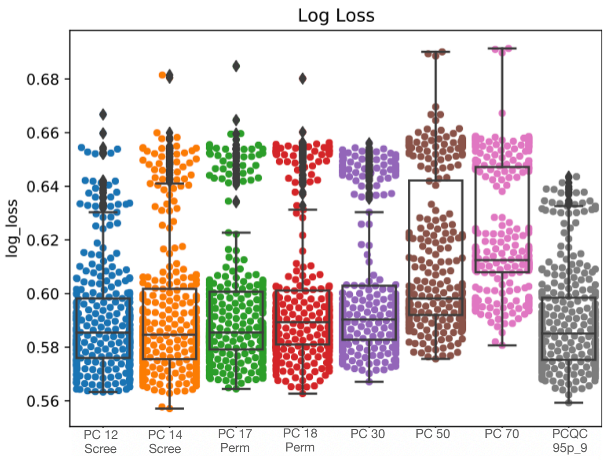


**Figure 17:** Log loss distribution for the 17k+ PBMC dataset for discerning between singlet and doublet cells.
